# Hands-free control of heterologous gene expression in batch cultures

**DOI:** 10.1101/150375

**Authors:** Olivier Borkowski, Drew Endy, Pakpoom Subsoontorn

**Author notes:** equally contributing authors (Olivier Borkowski, Pakpoom Subsoontorn). Email addresses: OB, DE, PS.

## Abstract

**Background:** Autonomous cell-based control of heterologous gene expression can simplify batch-culture bioprocessing by eliminating external monitoring and extrinsic control of culture conditions. Existing approaches use auto-induction media, synthetic cell-cell communication systems, or application-specific biosensors. A simpler, resource-efficient, and general-purpose expression control system responsive to common changes during batch culture would be useful.

**Results:** We used native *E.coli* promoters and recombinase-based switches to repurpose endogenous transcription signals for control of heterologous gene expression. Specifically, natural changes in transcription from endogenous promoters result in recombinase expression at different phases of batch culture. So-expressed recombinases invert a constitutive promoter regulating expression of arbitrary heterologous genes. We realized reversible and single-use switching, reduced static and dynamic cell-to-cell variation, and overall expression amplification. We used “off-the-shelf” genetic parts and abstraction-based composition frameworks to realize reliable forward engineering of our synthetic genetic systems.

**Conclusion:** We engineered autonomous control systems for regulating heterologous gene expression. Our system uses generic endogenous promoters to sense and control heterologous expression during growth-phase transitions. Our system does not require specialized auto-induction media, production or activation of quorum sensing, or the development of application-specific biosensors. Cells programmed to control themselves could simplify existing bioprocess operations and enable the development of more powerful synthetic genetic systems.

## Background

Unregulated expression of heterologous genes can hinder cell growth, reduce bioprocess yields, and increase the chance of selecting for mutants that disrupt system performance [1][2][3][4][5]. Regulated expression via chemically inducible promoters mitigates these problems by offering control of gene expression levels and timing [6]. For example, protein production processes often repress heterologous gene expression until high cell densities are reached, whereupon added inducers trigger expression (Figure 1A). However, the use of inducible promoters requires close monitoring of culture conditions so that extrinsic control signals are applied correctly.

**Figure 1:**
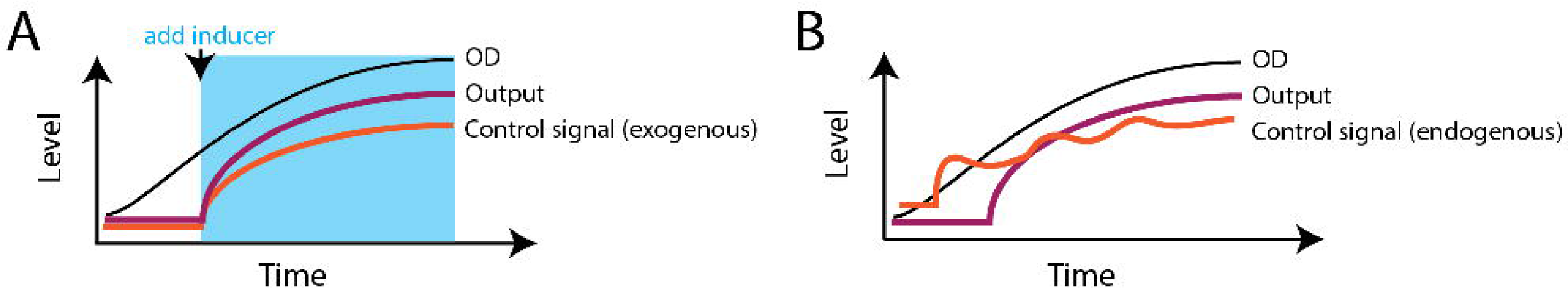
An autonomous switch could obviate the need to induce batch-culture production. (**A**) Schematic plot of a conventional heterologous protein expression exogenously controlled under a chemically inducible promoter. As batch-culture cell density (black line) reaches a target level, an experimenter adds a chemical inducer (shaded blue area). The inducer activates an exogenous transcription control signal (orange line) from a chemically inducible promoter that up-regulates a heterologous output gene (purple line). (**B**) Schematic plot of heterologous protein expression endogenously controlled under an autonomous switch. As cell density increases, growth-phase-responsive endogenous transcription signal increases (orange line) and eventually turns ON a heterologous output gene (purple line); no chemical inducer is needed.

Auto-induction media avoids the need for external monitoring of cell density by using catabolic repression to independently control induction timing [7][8]. In more detail, auto-induction media typically contains both an inducer (e.g., IPTG) plus a carbon source (e.g., glucose) or only a carbon source. The carbon source represses heterologous gene expression early in batch culture. Catabolite repression is relieved as the media is depleted due to cell growth resulting in late-phase heterologous gene expression. Natural quorum sensing systems have also been repurposed to control heterologous gene expression in a similar manner [9][10][11]. Specifically, cell-cell signaling molecules are produced by cells and secreted into the media (e.g., acyl homoserine lactones). Heterelogous gene expression is only activated only once the quorum signaling molecules accumulate above a threshold concentration.

Both auto-induction and synthetic quorum sensing rely on specific changes in the media to control expression across an entire culture. Another approach is to place heterologous gene directly under the control of endogenous signals within a cell [12]. For example, recent work by Dahl et al. [13] uses a farnesyl pyrophosphate (FPP) repressible promoter to control expression of a heterologous amorphadiene production pathway. In this case, FPP-mediated repression is used to mitigate harmful accumulation of FPP as a toxic intermediate metabolite. Similar projects have realized autonomous gene expression control by developing genetically encoded sensors for application-specific signals, which are then used to control heterologous gene expression on a case-by-base basis [14][15][16][17] (Table S1).

Auto-induction media has been widely used in laboratory-scale protein productions. However, inducer costs can prohibit their application in high volume fermentations. Moreover, the use glucose to repress early-phase expression can result in acetate levels that inhibit cell growth[18]. Synthetic quorum sensing systems eliminate the requirement for specialized media. But, so-engineered cells must actively produce and power the quorum sensing system itself in addition to the desired heterologous system [19]. Application-specific control systems can avoid both issues, but require ad hoc development of a purpose-built sensor and control system for each new pathway or project.

Here, we developed growth-phase dependent autonomous switches controlled by endogenous signals (Figure 1B). Our control systems do not require specialized media and use “non-volatile” digital genetic switches [20] [21] to minimize consumption of cell resources otherwise needed for growth and heterologous expression.

## Results

An ideal autonomous growth phase-responsive expression control system should work in typical culture conditions and realize performance latencies and dynamic ranges that match or improve upon existing expression control methods. Additionally, the system itself should consume minimal cellular resources, avoid specialized media requirements, and support use with any application.

### Selection of growth-phase-responsive endogenous transcription sources

We choose to explore the use of native promoters responsive to changes typical of batch culture as potential controllers of heterologous gene expression. For example, hundreds of native *E. coli* promoters have been shown to change in activity during entry into stationary phase [22][23][24][25]. To determine which endogenous transcription sources might be appropriate for use in reliably controlling heterologous gene expression we re-characterized native promoters, searching for promoters with the greatest change in expression levels, the least cell-cell heterogeneity, and minimal overlap between exponential and stationary phase expression levels. We also sought promoters whose specific activity on semi-solid media (i.e., plate-based overnight colonies) was lower than in exponential-phase liquid culture. These requirements, taken together, were formulated to help us realize engineered expression control systems having low latencies, high dynamic ranges, minimal spontaneous activation rates, and practical ease of use (below).

We used a synthetic promoter, P*j23101*, as a standard reference promoter for comparing the relative activities of candidate promoters on semisolid media and in exponential and stationary phase liquid cultures (Figure 2). We also re-characterized the frequently used P*tet* and P*bad* inducible promoters to help guide selection of suitable native promoters. We used a standardized green fluorescent protein (GFP) expression cassette and the relative promoter unit (RPU) framework developed by Kelly et al. to regularize quantitative comparison of promoter activities across conditions and experiments [26]. Both the level and activity of RpoS, the *E. coli* stationary-phase sigma factor, increases during entry into stationary phase [27]. Thus, we chose the promoter of *rpoS* as a candidate for our stationary-phase sensor. Previous work by Shimada et al. characterized rpoS-dependent stationationary-phase promoters [23]. Shimada et al. compared batch-culture GFP fluorescence as driven by each tested promoter in relation to a reference RFP fluorescence driven by a common constitutive promoter. The promoters for genes *gadA*, *dps*, *yiaG* and *hdeA* were observed to produce the highest GFP/RFP expression fold change from exponential to stationary phase. Therefore, we included these four promoters as our sensor candidates. For added context: GadA, a glutamate decarboxylase enzyme, is part of glutamate-dependent acid resistance system which confers resistance to extreme acid conditions [28]; Dps is a non-specific DNA binding protein that can form protein-DNA crystal to prevent DNA damages under various stresses [29]; YiaG is a predicted transcription regulator of unknown function (Uniprot); and, HdeA is a periplasmic protein important for acid stress responses [30]. We selected two additional promoter candidates taken from a gene *fumA* and *rpoA*. FumA is one of three fumerase isozymes in TCA cycle [31]; we selected the *fumA* promoter because its mRNA level is known to increase as growth rate decreases [32]. *RpoA* is RNA polymerase alpha subunit; RNA polymerase concentration decreases as growth rate decreases [33][34] and we expected downregulation of the *rpoA* promoter in stationary phase.

**Figure 2:**
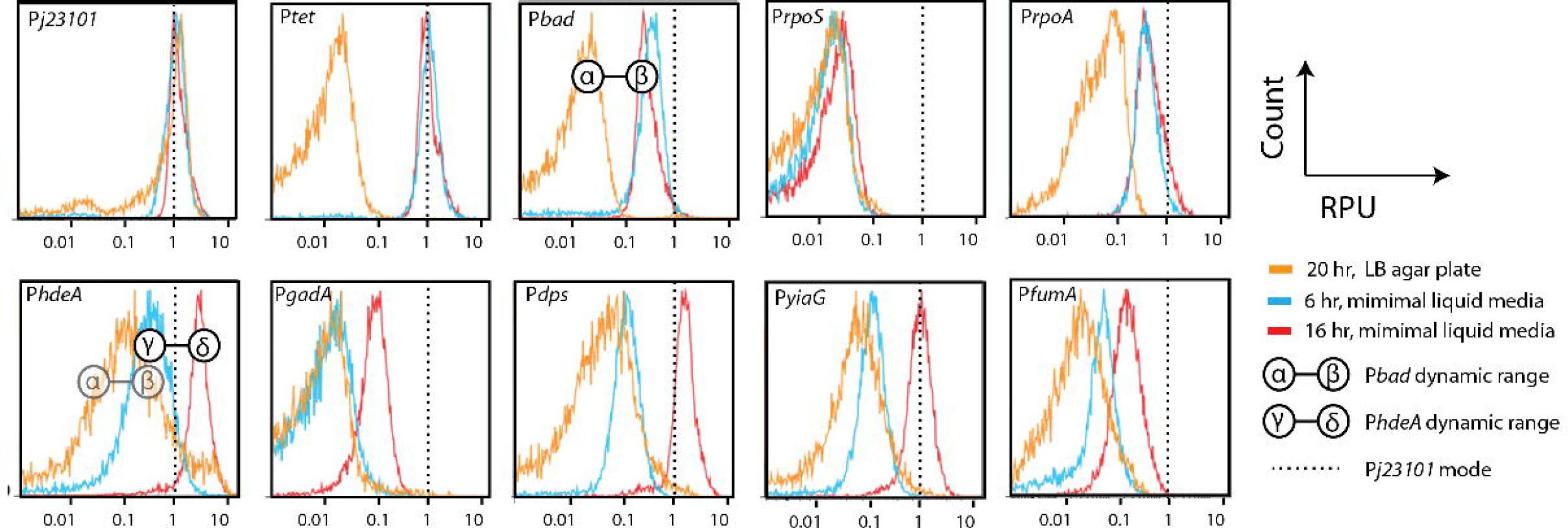
The *hdeA* promoter generates a dynamic range comparable to conventional inducible promoters. Each subplot shows single-cell strength distribution (in Relative Promoter Unit, RPU) of each tested promoter. Dashed lines mark the expression mode of the reference promoter P*J23101* which we used for normalizing presented promoter strength. We measured promoter strengths from colonies grown overnight on LB agar plate (orange), in liquid minimal media during exponential phase (blue) and during stationary phase (red). For P*tet* and P*bad* characterization, we had no inducer on LB agar plate but we added 20 ng/ml aTc and 0.1% arabinose respectively at the beginning of liquid culture. The α-β symbol marks the dynamic range of P*bad* before and after induction; the γ-δ symbol marks the dynamic range of P*hdeA* from exponential to stationary phase.

In previous studies, promoter activity measurements relied on using each promoter’s natural 5’ UTR directly coupled to a reporter gene open reading frame [23]. Here, we insulated our candidate promoters (Table S2) and via a standardized 5’UTR consisting of a self-cleaving ribozyme [35] and a bicistronic translation initiation sequence [36]. This standardized 5’UTR should reduce post-transcriptional variability of GFP expression and thus allow GFP measurements to better represent promoter activities.

Our reference promoter, P*j23101*, produces nearly constant GFP levels across three different growth conditions: overnight on solid LB agar, exponential liquid phase, and stationary liquid phase (Figure 2). We used the mode of the P*j23101*-driven GFP level to convert all measured GFP levels from arbitrary units to a Relative Promoter Unit (RPU) [26]. The *rpoS* and rpoA promoters also produced nearly constant expression levels across different growth conditions. Expression from the fumA promoter increased only slightly upon entering stationary phase. We found the *hdeA* promoter produced an average 20-fold change similar to the P*bad* promoter, albeit spanning a higher relative dynamic range but with almost entirely non-overlapping HIGH/LOW distributions at a single cell level. Since we had previously demonstrated functional recombinase switches controlled via P*bad* [20], we expected the native dynamic range of P*hdeA* activity to be similarly sufficient. However, the high basal level of P*hdeA* suggested that previous recombinase switches would spontaneously activate in response to P*hdeA-*initiated transcription, regardless of growth phase. Other tested promoters, P*gadA*, P*dps* and P*yiaG*, were also up-regulated during stationary phase but to a lesser degree than P*hdeA*.

We also observed the activity of our candidate stationary phase promoters from colonies grown overnight on solid media, and found expression levels to be as low as in exponential liquid culture (Figure 2: P*hdeA*, P*dps*, P*gadA* and P*yiaG*, orange and blue plots). Cloning process typically include plating transformant colonies on solid media followed by overnight growth. Therefore, having low promoter activity on solid media is important for the construction process of an autonomous growth-phase switch. Stated differently, if promoter activities are too high on solid media, the promoters might trigger state switching even before a liquid culture is innoculated.

### Functional composition of autonomous single-use switches

We constructed a single-use switch using a stationary-phase promoter to control an irreversible recombinase switch (Figure 3A). Specifically, a stationary-phase promoter controls expression of bacteriophage Bxb1 integrase; a constitutive promoter is located between opposing bacteriophage Bxb1 attB/attP recombination sites and oriented in the opposite direction from an output gene of interest (BP state). During growth-phase transition the stationary-phase promoter drives integrase expression which SETS the switch by recombining attB and attP site into attL and attR site (LR state). Consequently, the output promoter inverts toward the output gene of interest and turns the output gene ON. The irreversibility of integrase-mediated inversion maintains the output in an ON state even if cells are returned to exponential phase.

**Figure 3:**
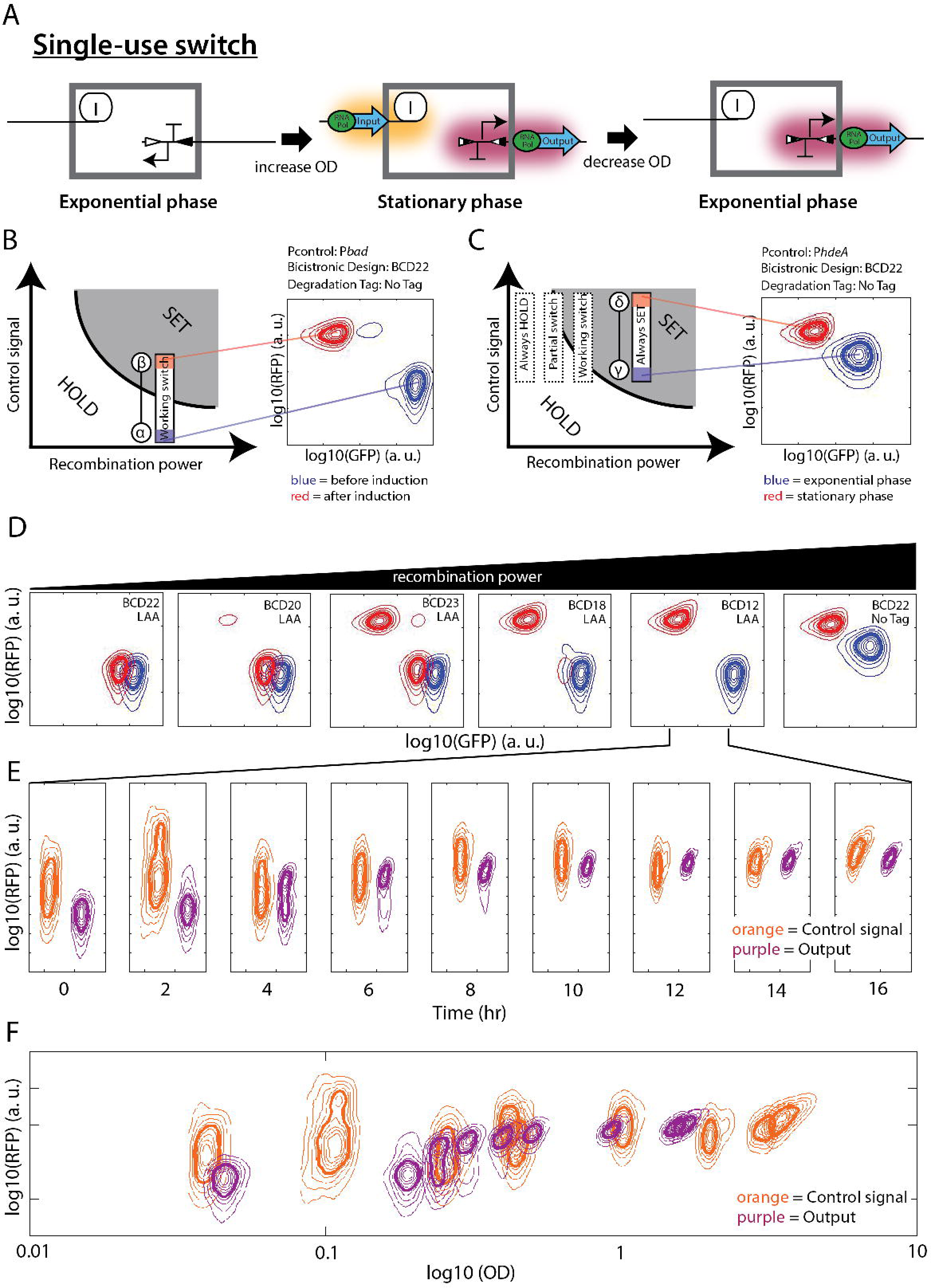
Functional composition and operation of an autonomous single-use switch. (**A**) Switch mechanisms. Hollow and solid triangles represent attB and attP sites; semi-solid triangles represent attL and attR sites. Glows mark transcriptionally active DNA. Green ellipses and blue arrows represent RNA polymerase and transcription current, respectively. (**B**) (Left) Schematic phase-diagram of switch behaviors with respect to control signal strength and recombination power. The areas labeled “SET” and “HOLD” represent parameter ranges that turn ON the switch and that keep the switch in OFF state, respectively. The rectangle represents the parameter range accessible to a functional arabinose inducible switch. The control signal strength without (blue end) and with (red end) arabinose induction falls within HOLD and SET region, respectively. (Right) Flow cytometry contours showing fluorescent distribution of the arabinose inducible switch without (blue) and with (red) arabinose, respectively. Thin contour interval encompasses 10% of population; thick contour encompasses 50% of population. (**C**) Schematic phase-diagram / flow cytometry contour similar to (B) but the switch is driven by a stationary phase promoter (P*hdeA*) instead of P*bad*. The solid rectangle represents the parameter range accessible to this switch. Dashed rectangles represent parameter ranges accessible to other switch variants with lower recombination powers. (**D**) Cytometry result similar to (C) (right) showing fluorescent output from P*hdeA* composed to integrase cassettes with different recombination powers. Upper right corner of each subplot notes translation initiation sequence and protein degradation tag used on the integrase cassette. (**E**) Output fluorescent distribution (purple) from a functional single-use switch responding to a control signal (orange) from P*hdeA.* Contours of switch output were shifted to the right to simplify the display. (**F**) The same data shown in (E) but plotted as functions of cell optical density (OD) instead of time. The center of the contours from each sample is placed horizontally at its corresponding sample’s OD. The width of each contour represents forward scatter.

From our previous work [21], we expected that applying a single-use recombinase switch as a signal relay between a stationary-phase promoter and an output gene should have two benefits. First, the recombinase switch should convert a continuously variable and potentially noisy transcription input signal from the stationary phase promoter to a stepwise transcription signal driving expression of the output gene. Such transcription signal “digitization” should occur because the invertible output promoter within the recombinase switch can exist in only two possible orientations, either OFF or ON. Second, the recombinase switch should increase fold-change of its relayed signal. Such transcription signal “amplification” occurs because transcription initiation from the output promoter is highly directional and a relatively small amount of recombinase is needed to realize switching. Thus, a small increase in control signal from the native growth-phase responsive promoter should produce enough integrase to flip the output promoter, resulting in large changes in the output signal. Of note, one drawback of a single-use switch is that any spontaneous switching failure is permanent. Specifically, any random bursts of integrase expression any time after the DNA encoding the autonomous switch is introduced in the cells would result in prematurely and irreversible activation of any heterologous genes controlled by the switch’s output.

The activity of the stationary-phase promoter and sensitivity of the recombinase switch together determine the output state of the autonomous single-use switch. The sensor determines the “control signal,” the number of RNA polymerase molecules transcribing the integrase gene at any given time. The level of transcription current depends on the choice of promoter and culture conditions. The recombinase switch’s sensitivity determines all post-transcription control of integrase expression. For convenience, we defined “recombination power” as a way of quantifying the propensity of any given recombinase switch to SET. For example, recombination power is high when integrase synthesis rates are high or integrase degradation rates are low. Together, the strength of the control signal from the growth-phase responsive promoter combined with the recombination power of a given recombinase switch determine the overall SET propensity for any given sensor/recombinase switch composition (Figure 3B).

We first built a chromosomally-integrated arabinose inducible switch using P*bad* as a transcription source and BCD22 as a weak translation initiation sequence (Table S3). An invertible constitutive promoter expresses GFP in the BP state (OFF) and RFP in the LR state (ON; Additional File 1: Figure S1). We demonstrated that the arabinose inducible switch was functional: at least 90% of cells were in an GFP expressing state in the absence of arabinose and more than 90% switched to an RFP expressing state after arabinose induction (Figure 3B, right). The stationary-phase promoter, P*hdeA*, has basal expression level on solid media and during exponential phase comparable to induced expression level from P*bad* (RPU ~ 0.2) (Figure 2). Thus, as expected, a P*hdeA*-driven integrase translated via BCD22 always switched into an RFP expressing state even prior to inoculation of liquid cultures (Figure 3C).

An integrase cassette with even the weakest bicistronic translation initiation sequence (BCD22) had too high recombination power for the P*hdeA* dynamic range. To further lower switch recombination power we added a wild-type ssrA degradation tag (noted as LAA in Figure 3D) at the C-terminus of the integrase to increase protein degradation rates [37]. We found that this new design could not switch to an RFP expressing state even after an overnight growth in liquid media, implying that recombination power was too weak to be responsive to P*hdeA* (Figure 3D, left most subplots). We then changed the bicistronic translation initiation sequence of the integrase to a rational series of stronger off-the-shelf designs (BCD20, BCD23, BCD18 and BCD12). The switch using BCD12 functioned properly (i.e., remaining in a low RFP/high GFP state at the beginning of the culture and switching completely to a high RFP/low GFP state at the end of the culture (Figure 3D, box 5)). We observed incomplete state switching when using BCD20, BCD23 and BCD18 (Figure 3D, boxes 2, 3 and 4). The rank order of switching efficiencies closely matched the rank order of known BCD strengths as measured in a prior study [36]. We also showed that the rank order of switching efficiency was preserved using a different stationary-phase promoter, P*dps* (Additional file 1:Figure S2).

We next tested whether using a recombinase switch as a signal relay improves the quality of the transcription signal relative to that of the stationary-phase promoter (P*hdeA*). We compared RFP time-course kinetics driven directly by P*hdeA* to RFP time-course kinetics driven through a single-use autonomous switch (Figure 3E and 3F, orange and purple, respectively). Overall, both P*hdeA*-mediated and switch output expression levels increased from the beginning of the liquid culture to stationary phase 16 hr later. However, the direct control signal from P*hdeA* had higher cell-to-cell variability, especially during earlier time points in liquid culture. Moreover, while the relayed output monotonously increased, the direct control signal from P*hdeA* varied over time; this fluctuation was consistent across repeated experiments (Additional file 1:Figure S3). The stationary-phase promoter P*hdeA* is known to be controlled by a complex natural regulatory network (Figure S7) [38][39] which presumably enables individual cells to fine-tune *hdeA* expression to specific physiological states. The use of a recombinase switch as a relay had at least two effects. First, the recombinase switch transposes transcription output from P*hdeA* to transcription output from a constitutive promoter, P*j23119*, located between recombination sites. Since P*j23119* is unregulated, it is possibly less responsive to cell-to-cell variability. Second, the recombinase switch functions as a signal integrator at a population level; above a certain threshold switch output turns ON irreversibly. Therefore, the amount of ON state cells always increases even if the control signal fluctuates. Stated differently, the time-course integral of any fluctuating input control signal always results in a monotonic increase in output signal. We were further able to rationally tune the kinetics of switch output using integrase cassettes with different recombination powers, with higher recombination power producing faster switching speeds, as expected (Additional file 1:Figure S4).

### Functional composition of autonomous reversible switches

We also developed autonomous reversible switches based on our previously engineered rewritable recombinase switch [20]. The reversible system consists of a growth-phase-independent integrase cassette plus a growth-phase-responsive excisionase-integrase bicistron; both cassettes are place together on a high copy plasmid. An output promoter (P*j23119*) is again located between opposing attL/attR recombination sites and oriented in the opposite direction from an output gene on the genome (Figure 4A). When the culture enters stationary phase, the stationary-phase promoter drives expression of excisionase and integrase, recombining attL/attR sites into attB/attP sites. Consequently, the output promoter inverts toward the output gene and turns the output gene ON. If the culture returns to an exponential phase, excisionase levels decrease and the presence of integrase alone converts attB/attP back to attL/attR, turning OFF the output gene (Figure 4A).

**Figure 4:**
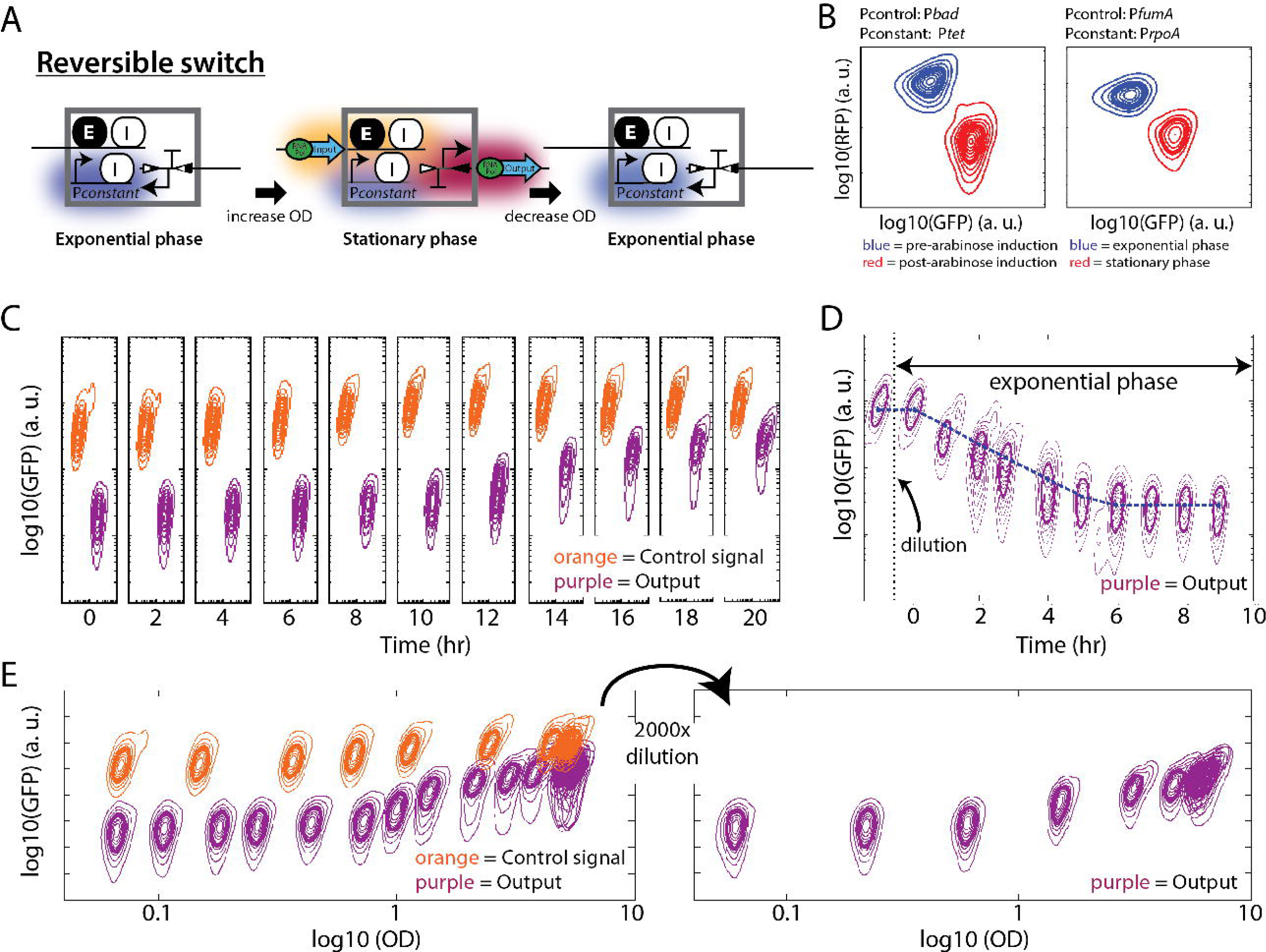
Functional composition and operation of an autonomous reversible switch. (**A**) Switch mechanisms. Hollow and solid triangles represent attB and attP sites; semi-solid triangles represent attL and attR sites. Glows mark transcriptionally active DNA. (**B**) (Left) Fluorescent distribution of a functional ara/aTc inducible switch [18] in liquid culture containing aTc without (blue) or with (red) arabinose. Thin contour interval encompasses 10% of population; thick contour encompasses 50% of the population. (**B**) (Right) Fluorescent distribution of a functional autonomous reversible switch with P*rpoA* driven integrase and P*fumA* driven excisionase-integrase bicistron during an exponential phase (blue) and a stationary phase (red). (**C**) Fluorescent distribution of a functional autonomous reversible switch outputs (purple) responding to a control signal from a control promoter, P*fumA* (orange). Contours of switch output were shifted to the right to simplify the display. (**D**) Fluorescent distribution of an autonomous reversible switch returning from an ON state in a stationary phase to an OFF state in an exponential phase. After 100x dilution, we kept the cells in an exponential phase by repeated dilutions. The dashed line represents the predicted decreasing GFP level due to dilution from cell growth alone. (**E**) (Left) The same data shown in (C) but plotted as a function of optical density. (**E**) (Right) Fluorescence measurements from an autonomous reversible switch output after dilution from a saturated phase into fresh media. In an exponential phase, cells reversed from an ON state to an OFF state and autonomously switched to an ON state again during the transition to a stationary phase. The center of the contours from each sample is placed horizontally at its corresponding sample’s OD. The width of each contour represents forward scatter.

We experimented with different transcription sources to control Bxb1 integrase and excisionase in our autonomous reversible switch. Similar to our single-use switch, a reversible switch had an invertible promoter drive RFP expression in LR state (OFF) and GFP expression in BP state (ON, Additional file 1: Figure S1). We started from our original rewritable recombinase switch that used P*tet* and P*bad* to drive integrase and integrase-excisionase bicistron, respectively (Figure 4B, left) [20]. We first replaced P*bad* with a stationary-phase promoter, P*hdeA*, as a transcription source of the integrase-excisionase bicistron cassette; the integrase cassette remained under P*tet* control. This construct did not function properly: cell populations remained in an intermediate state expressing both GFP and RFP in both exponential and stationary phases (Additional file 1: Figure S5 A, second box). We hypothesized that integrase expression levels from uninduced P*tet* were not high enough to catalyze complete state switching. We increased integrase expression levels by replacing P*tet* with P*rpoA.* However, this design remained in an RFP expressing state in both exponential and stationary phase (Additional file 1: Figure S5 A, third box). We hypothesized that the level of integrase expressed from P*rpoA* was too high compared relatively to the level of excisionase expressed from P*hdeA* to allow for LR to BP state switching. We then tried replacing the growth-phase promoter P*hdeA* with P*fumA* (Figure 2). This design functioned properly; cells remained in RFP expressing state during exponential phase and switched to a GFP expressing state during stationary phase (Figure 4B, right).

We compared a direct control signal from P*fumA* (Figure 4C, orange contour) to a relayed signal passed through a reversible switch (Figure 4C, purple contour). We found that GFP expression driven by P*fumA* directly increased 2.5-fold and reached its maximum level after 10 hours; GFP expression driven by the relayed signal increased 14-fold and reached its maximum level after 18 hours (Figure 4C, additional file 1: Figure S5). Stated differently, the reversible switch amplified expression fold change (~5.6 fold) but also caused an 8 hour expression delay. Quantitative PCR results showed that DNA inversion of the reversible switch occurred by 10 hours, implying that the apparent switching delay results from transcription-translation delay of GFP reporter and not from DNA inversion kinetics (Additional file 1:Figure S6 A). We could reverse an ON state switch to an OFF state switch by diluting the culture back to exponential phase in fresh medium. GFP expression returns to an OFF state level within 4-5 hours following dilution (Figure 4D). However, quantitative PCR results showed that DNA reversion was complete within less than one hour post dilution (Additional file 1:Figure S6 B). GFP dilution due to cell growth alone can explain the observed loss of GFP after the switch returns to the OFF state (Figure 4D, blue dashed line). Following state reversion, the reversible switch could autonomously turn ON again as a culture reenters stationary phase (Figure 4E, Additional file 1:Figure S6 C).

## Discussion

We show that digital amplifying genetic switches can be reliably used for autonomous control of gene expression. We engineered reversible and irreversible recombinase switches to relay transcription signal between a stationary phase promoter and any heterologous gene. By using a stationary phase signal our designs overcome limits faced by previous autonomous systems that need specialized media, complex synthetic circuits, or application-specific biosensors.

The development of our autonomous switches confirmed that the recombinase-based switch constructed by Bonnet et al. is easy to manipulate and robust to modifications. Specifically, we used well characterised, standardized parts including recombinase-based switches, bicistronic translation initiation elements, ribozymes and protein degradation tags to rationally and reliably tune recombinase expression and activity within just a few-test iterations. We realized easy modification of the promoter controlling recombinase expression and expression level tuning. Likewise, the modularity of our systems allowed modifying the strength of the output signal without altering the recombinase expression level. Finally, the switches filter and reduce input signal fluctuations and produce an amplified output signal, which taken together should reduce cell-cell heterogeneity in industrial scale fermentations.

We provide simple yet efficient tools for batch culture bioproduction with comparable expression fold change and kinetics as previously obtained via autoinduction media. Our switches should be useful for optimizing metabolic pathway performance by reducing the burden of heterologous expression on the host, as the switches decouple growth phase and protein production. Autonomous genetic switches controlled by carefully selected endogenous signals should find many uses, from affordable and reliable control of industrial scale fermentations to selective control of gene expression in response to changing environmental conditions (e.g., to help prevent escape of engineered strains from one environment to another).

## Materials and Methods

### Plasmids and strains

Plasmids were transformed using standard procedures [48] in chemically competent *E. coli* DH5alphaZ1 and plated on LB agar plates containing the appropriate antibiotics. Plasmids were constructed using standard BioBrick™ [49], Gibson assembly [50] or Golden Gate assembly [51]. Coding sequences for Bxb1 integrase and excisionase were synthesized by DNA 2.0 (Menlo Park, CA, USA).

Parts encoding P*bad* (iGEM registry accession number: BBa_I0500), superfolder GFP (iGEM registry accession number: BBa_I1746916), mKate2 [52], and the P*tet* (BBa_R0040) were obtained from the iGEM Registry of Standard Biological Parts (http://parts.igem.org/). Growth-phase-dependent transcription initiation sequences were amplified by PCR from *E. coli* DH5alphaZ1 genome. Since the boundaries of some growth-phase-dependent transcription initiation sequences are unknown, we simply took 400 (P*fumA* and P*rpoA*), 500 (P*rpoA*, P*hdeA*, P*gadA*, P*dps* and P*yiaG*) or 754 (P*rpoS*) bases upstream of the start codon of growth-phase-responsive-genes and used them as growth-phase-responsive transcription sources (Table S3).

PCR reactions were performed using the platinum Hi-Fi PCR supermix (LifeTechnology, USA) and Phusion Hi-Fidelity DNA polymerase Mix (Biolabs USA), using a 1 min extension time per kilobases. Primers were purchased from IDT.

The recombination target of the switch, called “Flippee Targets,” (Additional file 1:Figure S1) consists of the constitutive promoter P*j23119* flanked by opposing attB and attP sites (BP states) or flanked by opposing attB and attP sites (LR states). The BP and LR states drive the expression of GFP and mkate2, respectively. The Flippee Targets were first cloned on a low copy plasmid (pSB4A5) then integrated into *E. coli* DH5alphaZ1 chromosome using Phi80 phage integration sites [53].

For the single-use switch, the growth-phase-responsive promoter drives Bxb1 integrase with RiboJ and different BCD bicistronic junctions at 5’UTR as indicated in the main text (Additional file 1:Figure S9). This design is integrated in the genome at the HK022 sites.

For the reversible switch, the control promoter drives Bxb1 integrase and excisionase with strong RBS-50000 [45] and ARH degradation tag (Additional file 1:Figure S10). The constant promoter P*rpoA* drives Bxb1 integrase with a randomized RBS 6N. This design is on J64100 plasmid.

### Characterization of growth-phase-responsive promoters

Constructs with a growth phase responsive promoter driving GFP were transformed in chemically competent *E. coli* on LB plate containing 5 ug/ml chloramphenicol and then grown overnight at 37°C. For LB agar plate measurements, several colonies were picked and resuspended in 200 μl 60% glycerol for later FACS measurements. For measurements in minimal liquid media, a single colony of *E. coli* was grown overnight in 1 ml minimum media (Hi-Def Azure media, Teknova) + 0.6% glycerol with 5 ug/ml chloramphenicol on a 96 wells plate at 37°C, 480 rpm, 80% humidity. Then, fresh minimal media was added to adjust each overnight samples to OD=1. Adjusted overnight samples were diluted 1:1000 in 1 ml minimal media with 5 ug/ml chloramphenicol and grown on a 96 wells plate at 37°C, 480 rpm, 80% humidity. After 5 hours, 200 μl sample was taken from the 96 wells plate and added to 200 μl 60% glycerol solution and stored at −80°C freezer. After 16 hours, another 200 μl sample was taken and stored as glycerol stock at −80°C freezer. Before FACS measurement, 100 μl of the 5 hour and 16 hour samples were thawed on ice and added to 1 ml of sterile water in FACS tube. Flow cytometry analysis was performed using LSR II cytometer (BD-Bioscience) at the Stanford Shared Facs Facility (SSFF) with the following parameters: threshold at 800 V SSC, Laser 603 V FSC, 350 V SSC, 560 FIT-C and 650 PE-Texas-RED.

### Switches characterization

Single-use and reversible switches were transformed in chemically competent *E. coli* on LB plate containing 5 μg/ml chloramphenicol. After overnight (20 hours) at 37°C, colonies on the plate were resuspended together in LB media. Final OD was adjusted to approximately 2-3 and stored as glycerol stocks. To start a liquid culture, we scraped the glycerol stock with 1 ml micropipet tip, inoculated in 1 ml Hi-Fef Azure media +0.6 % glycerol and 5 ug/ml chloramphenicol and grew at 37°C, 480 rpm, 80% humidity. The exponential phase and the stationary phase samples were taken at 6 hour and 16 hour, respectively, after inoculation. For chemically inducible promoters (P*tet* and P*bad*), inducers were added to the media (20 ng/ml aTc and 0.1% arabinose, respectively) at the beginning of liquid cultures.

### Time-course experiment of single-use autonomous switch

The glycerol stocks for a strain with P*hdeA* driven mKate2 and a strain with P*hdeA*driven single-use switches were prepared as described in the previous section. To start a liquid culture for P*hdeA* time course experiment, 50 μl glycerol stock aliquot of P*hdeA*-mKate2 was inoculated in 10 ml High-Def Azure media + 0.6% glycerol 5 μg/ml chloramphenicol and grown at 37°C, 480 rpm, 80% humidity. To start a liquid culture of P*hdeA*-driven single-use switches, 100 ul glycerol stock aliquot of P*hdeA*-mKate2 was inoculated in 10 ml High-Def Azure media +0.6% glycerol 5 μg/ml chloramphenicol 10 ug/ml kanamycin and grown at 37°C 480 rpm, 80% humidity. We took 1 ml sample every two hours and stored as glycerol stock for FACS measurement.

### Time-course experiment of a reversible autonomous switch

The glycerol stocks for a strain with P*fumA* driven superfolder GFP and a strain with P*fumA*-driven reversible switch were prepared as described in the previous section. To start a liquid culture for P*fumA* time course experiment, 50 μl glycerol stock aliquot of P*fumA*-GFP was inoculated in 10 ml High-Def Azure media + 0.6% glycerol 5 ul/ml chloramphenicol and grown at 37°C, 480 rpm, 80% humidity, overnight. To start a liquid culture of P*fumA*-driven reversible switch, 100 μl glycerol stock aliquot was inoculated in 10 ml High-Def Azure media + 0.6% glycerol, 5 μg/ml chloramphenicol 10 ug/ml kanamycin and grown at 37°C, 480 rpm, 80% humidity. Overnight cultures were diluted 1:2000 in fresh media, a first 1 ml sample was taken after culture reaching OD=0.01 and every two hours 1 ml samples were stored as glycerol stock for FACS measurement.

### Quantitative PCR measurements

Colony PCR was performed for BP percentage measurements at the genome level in cell population containing reversible switch. For the colony PCR, we used the Mix KAPA2G Robust DNA Polymerase with dNTPs (Kapa Biosystems, USA). Each tube contained 50 μl of reaction with 0.5 μl of each primer 100mM, 25 μl of KAPA2G polymerase Mix, 24 μl of sterile water and the resuspended colony. Then, the PCR reactions were carried out in the S1000 thermal Cycler instrument (Bio-Rad, USA) starting with a denaturing step for 30 seconds at 98°C and followed by 40 cycles (98° C, 10s; 65° C, 1min; 72° C, 5min).

BP percentages were quantified with real-time qPCR using the Fast Plus EvaGreen qPCR Master mix (Biotum, USA). Each 96 well plates contained 50 μl of reaction with 0.5 μl of each primer 100 mM, 25 μl of qPCR Master mix, 23.5 μl of sterile water and 0.5 μl of colony amplified DNA diluted a 100 time. For each sample, two combinations of primers were used: (i) primers amplifying the sequence between the promoter and the *gfp* superfolder sequence (BP state): GTACTAATCGGCTTCAACGTGCCG and TGGTAGTGATCAGCCAGCTGC (ii) primers amplifying the sequence between the promoter and the *mkate2* sequence (LR state): GTACTAATCGGCTTCAACGTGCCG and GACAGTACCTTCCATATATAATTTCATGTGCATATTTTCTTTAATTAATTC The qPCR reactions were carried out in the Bio-Rad iCycler Optical Module (Bio-Rad, USA) starting with a denaturing step for 30 seconds at 95°C and followed by 40 cycles (95°C, 30s; 55°C, 30s; 72°C, 25s). The measured abundance of switch in each state is given relative to the standard reference switch concentration.

### Predicted protein dilution kinetic

We used ordinary differential equation (ODE) to simulate the dynamical behaviors of a protein concentration affected only by the dilution and compare it to the fluorescence measured with the reversible switch after regrowing back to exponential phase (Fig. 4D). The model only takes into account protein dilution (μ) and no production or degradation.

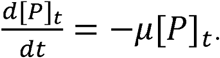

In other words,

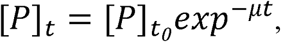

where μ is the growth rate (h^-1^), [P]_t_ is a measure of the GFP fluorescence level, [P]_t0_ is the measure of the GFP fluorescence level obtained immediately after dilution. To calculate μ we plotted the log(OD) versus time then estimated the slop of individual time point with respect to the previous and next time points. The OD was measured every 30 minutes over the span of 9 hours.

|  |  |  |  |  |  |  |  |  |  |  |
| --- | --- | --- | --- | --- | --- | --- | --- | --- | --- | --- |
| Time (h) | 0.5 | 1.0 | 1.5 | 2.0 | 2.5 | 3.0 | 3.5 | 4.0 | 4.5 | 5.0 |
| Growth rate ($\text{h}^{-1}$) | 0.52 | 0.52 | 0.55 | 0.64 | 0.61 | 0.63 | 0.61 | 0.63 | 0.60 | 0.60 |
| Time (h) | 5.5 | 6.0 | 6.5 | 7.0 | 7.5 | 8.0 | 8.5 |  |  |  |
| Growth rate ($\text{h}^{-1}$) | 0.64 | 0.61 | 0.60 | 0.60 | 0.60 | 0.62 | 0.62 | | | |

## Additional files

Additional file 1: Supplemental tables and figures.

Additional file 2: Construct Sequences.

## Authors’ contributions

PS and OB designed and conducted experiment. PS, OB and DE conceived of the study and wrote the manuscript. All authors read and approved the final manuscript.

## Acknowledgements

We thank P Jaschke, J Bonnet, and B Townshend for discussion. PS is supported by Stanford Bio-X Bioengineering Graduate Fellowship and Siebel scholar foundation. OB is supported by INRA CJS Fellowship. DE acknowledges generous support from NSF Synthetic Biology Engineering Research Center and Stanford University.

